# Enhancing lifespan of budding yeast by pharmacological lowering of amino acid pools

**DOI:** 10.1101/2020.10.30.362459

**Authors:** Nathaniel L. Hepowit, Jessica K. A. Macedo, Lyndsay E. A. Young, Ke Liu, Ramon C. Sun, Jason A. MacGurn, Robert C. Dickson

## Abstract

The increasing prevalence of age-related diseases and resulting healthcare insecurity and emotional burden require novel treatment approaches. Several promising strategies seek to limit nutrients and promote healthy aging. Unfortunately, the human desire to consume food means this strategy is not practical for most people but pharmacological approaches might be a viable alternative. We previously showed that myriocin, which impairs sphingolipid synthesis, increases lifespan in *Saccharomyces cerevisiae* by modulating signaling pathways including the target of rapamycin complex 1 (TORC1). Since TORC1 senses cellular amino acids, we analyses amino acid pools and identified 17 that are lowered by myriocin treatment. Studying the methionine transporter, Mup1, we found that newly synthesized Mup1 traffics to the plasma membrane and is stable for several hours but is inactive in drug-treated cells. Activity can be restored by adding phytosphingosine to culture medium thereby bypassing drug inhibition, thus confirming a sphingolipid requirement for Mup1 activity. Importantly, genetic analysis of myriocin-induced longevity revealed a requirement for the Gtr1/2 (mammalian Rags) and Vps34-Pib2 amino acid sensing pathways upstream of TORC1, consistent with a mechanism of action involving decreased amino acid availability. These studies demonstrate the feasibility of pharmacologically inducing a state resembling amino acid restriction to promote healthy aging.

## Introduction

Aging is the prime risk factor for fatal diseases including cardiovascular disease, diabetes, dementias and cancers [1–4]. Indeed, beyond 65 years of age the likelihood of Alzheimer’s disease doubles every five years with similar increases seen for other age-related diseases [5]. Strategies to slow the aging process and improve heath in older people are being investigated. Amongst these, nutrient restriction continues to gain support from studies in model organisms and clinical trials in humans as a way to promote healthy aging [6–12]. However, there is a critical need for alternative strategies to achieve nutrient restriction as few humans can limit nutrient intake to the degree required to improve health. Pharmacological approaches are a viable alternative to achieve nutrient restriction but finding compounds that induce the wide and complex array of cellular effects produced by nutrient restriction has been difficult [13]. Previously we found enhanced longevity in cells treated with myriocin (Myr, ISP-1) which targets the first and evolutionarily conserved step in sphingolipid synthesis [14]. Remarkably, as we show here, Myr lowers the free pool of most amino acids including ones known to enhance lifespan. These results pose the tantalizing question: How does slowing sphingolipid synthesis rewire metabolism and promote longevity?

Sphingolipids are abundant components of the plasma membrane (PM) in all eukaryotes. In the budding yeast *S. cerevisiae,* sphingolipids comprise 7-8% of the total mass and about 30% of phospholipids in the PM [15–17]. Myr was identified in screens for antibiotics [18] and rediscovered from a different source in a search for anti-inflammatory compounds [19]. Myr impairs activity of serine palmitoyltransferase (SPT) [20], the enzyme catalyzing the first step in sphingolipid synthesis which is ubiquitous in eukaryotes and vital for survival [21–23].

Previously we found a reduction in two of the three major sphingolipids in the yeast PM including inositol-phosphoceramide (IPC) and mannose-inositol-phosphoceramide (MIPC) (Figure 1A) in Myr-treated cells [14]. Remarkably, Myr treatment more than doubled chronological lifespan (CLS), a measure of how long yeast cells remain viable in stationary phase [14, 24]. Extensive transcriptional changes fostered by drug treatment overlapped significantly with those found in caloric restricted or rapamycin-treated long-lived yeasts. Myr treatment modulated a wide range of cellular processes with known roles in longevity including increased oxidative stress resistance, autophagy, genome stability, reduced ribosome and protein synthesis and rewired carbon, nitrogen and energy metabolism. Control of these processes required nutrient sensing pathways including the target of rapamycin complex I (TORC1), protein kinase A (PKA) and Snf1 (mammalian AMPK), all with established roles in longevity [1, 25, 26].

**Figure 1.**
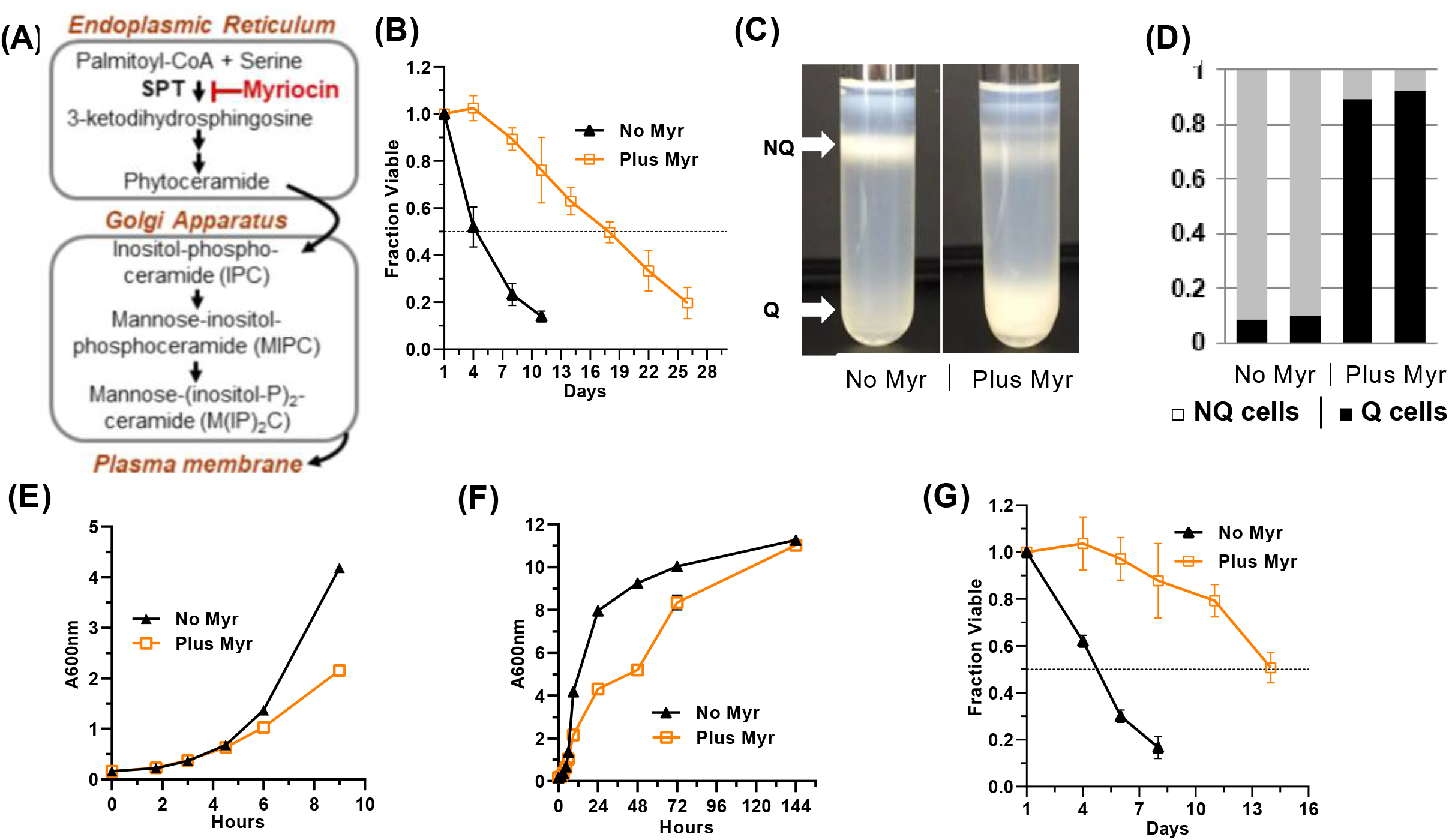
Effects of Myr on quiescence and lifespan. (A) Outline of the sphingolipid biosynthesis pathway in yeasts. (B) Myr treatment increases lifespan of BY4741 cells grown in 25 ml cultures. Statistical significance between drug-treated and untreated cells was determined by the Area Under Curve (AUC, 95% CI: 3.515-5.178 vs 14.77-18.05). Error bars: SEM. All data shown in this Figure were done with prototrophic BY4741 cells grown in SDC medium. (C) Representative density gradients showing the difference in the distribution of quiescent (Q) and non-quiescent (NQ) cells on day 3 of a lifespan assay in cultures without and with Myr treatment. (D) The number of cells in the Q and NQ bands from a density gradient were counted and plotted as a fraction of total cells. Data for two gradients of cell treated or not treated with Myr are shown. (E) Myr treatment begins to slow growth around 4 h after cells are inoculated into 200 mls of medium. (F) Myr treatment slows growth initially, but by 144 h the cell density of treated and untreated cells is similar. (G) Lifespan assay of cells grown in 200 ml cultures (AUC, 95% CI: 3.561-4.072 vs 9.962-12.78).

Although caloric restriction, a reduction in daily energy intake of up to 40%, has been and still is the major focus of many studies, other forms of nutrient restriction show potential for improving healthspan [4, 12]. Amongst these, protein and amino acid restriction promote health and longevity through multiple mechanisms to maintain normal physiological homeostasis [6–8, 27]. While lowering total protein intake fosters healthful effects, reducing intake of individual or groups of amino acids also improves health across species. Methionine (Met) restriction enhances longevity in organisms ranging from yeasts to worm, flies and mammals including rodents (mice and rats) plus the naked mole rat [28–33]. Reduction of branched-chain amino acids leucine, isoleucine and valine also promotes longevity in model organisms [27, 34]. A common feature of these interventions is to reduce TORC1 activity.

To uncover initial molecular mechanisms for Myr-induced longevity, we sought to understand how slowing sphingolipid synthesis by Myr treatment reconfigures cellular processes and reduces TORC1 activity. The TORC1 protein complex is a central regulator of longevity which senses multiple extra and intracellular signals and promotes appropriate responses to maintain cellular homeostasis [25, 26, 35]. Early studies of rapamycin and its inhibitory effect on TORC1 revealed a role in amino acid transport [36]. Numerous subsequent studies in yeasts established TORC1 as a key sensor of amino acids and other nitrogen sources [37, 38]. Together with our previous work, these studies hinted that Myr-induced remodeling of amino acid signaling or sensing might be part of the mechanism for increased lifespan.

Here we show that an early effect of Myr treatment is to maintain a low free pool of at least 17 amino acids in contrast to the expansion of pool size found in untreated cells. To understand the molecular basis of this effect, we investigated the trafficking and transport activity of Mup1, the major high affinity Met transporter in yeast. Although chronic exposure to Myr induced the endocytic clearance of Mup1, this clearance occurred much later than the observed drop in intracellular Met free pool, indicating that Myr treatment acutely inhibits transport function at the PM. These findings suggest that an acute effect of Myr treatment is diminished function of amino acid transporters (AATs) at the PM, resulting in lower amino acid availability in the cytosol that promotes longevity.

## Results

### Myriocin increases CLS in prototrophic yeast cells by inducing quiescence

Although our previous studies demonstrated that Myr extends the CLS of yeast, all of the data relied on auxotrophic or nutrient-requiring yeast strains [39]. Conceivably, Myr might only extend CLS in auxotrophic strains because it impairs uptake of one or more of the required nutrients thereby producing a state of nutrient restriction. Furthermore, overall amino acid homeostasis is perturbed in auxotrophic cells and masks how wild-type (WT), prototrophic cells respond to nutrient changes and environmental threats [40]. Thus, for the studies described here we used BY4741 cells made prototrophic with a plasmid complementing all mutant genes as was done for analysis of amino acid pools in the yeast deletion strain collection [41].

Indeed, we find statistically significance enhancement of CLS with untreated prototrophic BY4741 cells dropping to 0.5 fraction viable by day 4±1 while Myr-treated cell don’t drop to this level until day 14±2 (Figure 1B, Area Under the Curve, AUC, 95% CI: 3.515-5.178 vs 14.77-18.05). Lifespan assays with triplicate biological replicas we done 5 times with similar results. These findings validate previous CLS analysis of Myr-treated cells and reveal that the extended lifespan in the presence of Myr is also observed in prototrophic yeast strains.

The SDC culture medium used our current studies is weakly buffered with phosphate (pH 6) and some have maintained that lifespan enhancing strategies work in such conditions by protecting cells against acidification and accumulation of acetic acid [24]. We previously showed that Myr treatment enhanced lifespan of auxotrophic strains grown in SDC media buffered weakly or strongly, thus eliminating the possibility that drug treatment extended lifespan primarily by protecting cells against medium acidification [39]. We now find that Myr treatment increases lifespan in prototrophic BY4741 cells grown in strongly buffered SDC medium (Supplementary Figure 1). SDC culture medium has a relatively high concentration of amino acids, which increases lifespan by suppressing hyperacidification of culture media [42].

A hallmark of long-lived yeast strains is the ability to enter a quiescent state referred to as Q and characterized by the ability to survive in the absence of nutrients and then enter the cell division cycle when nutrients become available. In contrast, strains with a normal lifespan enter a senescent or non-quiescent (NQ) state typically found when cells cease growing and enter stationary phase where they rapidly loose viability [43, 44]. Yeast Q cells represent a physiological state similar to G0 cells in higher eukaryotes as exemplified by stem and other long-lived cell types [45].

To determine if Myr treatment promotes entry into the Q state, we analyzed cells at day 3 of a CLS assay and found that drug treatment induces more than 90% of prototrophic BY4741 cells to enter a Q state while less than 10% of untreated cells enter a Q state (Figures 1C and 1D). These data are consistent with our published data showing that Myr treatment reduces activity of TORC1, a highly conserved feature of longevity and entry into quiescence [45], and the activity of one of its effector pathways, the Sch9 kinase (mammalian S6K) [39]. Cells lacking Sch9 activity *(sch9*Δ cells) are long-lived and enter the Q state [46] as do cells undergoing nutrient limitation and reduction in TORC1 activity [44, 45, 47, 48].

### Free amino acid pools are smaller in Myr-treated cells

To understand how Myr reconfigures cellular physiology to enhance longevity, we optimized culture conditions to examine free intracellular amino acid pools (Materials and Methods). Under these conditions, 50-60% of cells bud after about 90 min of growth and complete their initial cell cycle after 2-3 h of growth (Supplemental Figure 2). The growth rate of Myr-treated cells slows at around 4±0.5 h of exposure (Figure 1E). Notably, Myr does not significantly reduce final cell density, since by day 1 of a CLS assay (72 h, Figure 1F) most cells have stopped growing with only a small, variable fraction of 0-20% slowly finishing their final cell cycle before entering quiescence on day 3 (144 h, Figure 1G).

To examine free intracellular amino acid pools, we first used a classical heat-extraction protocol followed by derivatization of amine groups with a fluorescent tag, compound separation by high-pressure liquid chromatography (HPLC) and quantification of fluorescent signals in eluted compounds. Stationary phase cells were inoculated into fresh culture medium and a sample was taken immediately and then at 1 h intervals. We find that the 18 amino acids detected by this procedure show a statistically significant difference in pool size with untreated cells having a larger pool than drug-treated cells (Figures 2A) (Cys not detectable and Ala not changed by Myr-treatment). In untreated cells, the pool of most amino acids has begun to increase by the 1 h time point. In contrast, most pools remain relatively constant in Myr-treated cells. This effect of Myr was independent of pool size and was observed for amino acids normally maintained at intracellular concentrations that are relatively low (e.g., Met and Tyr) or high (e.g., Glu and Ser).

**Figure 2.**
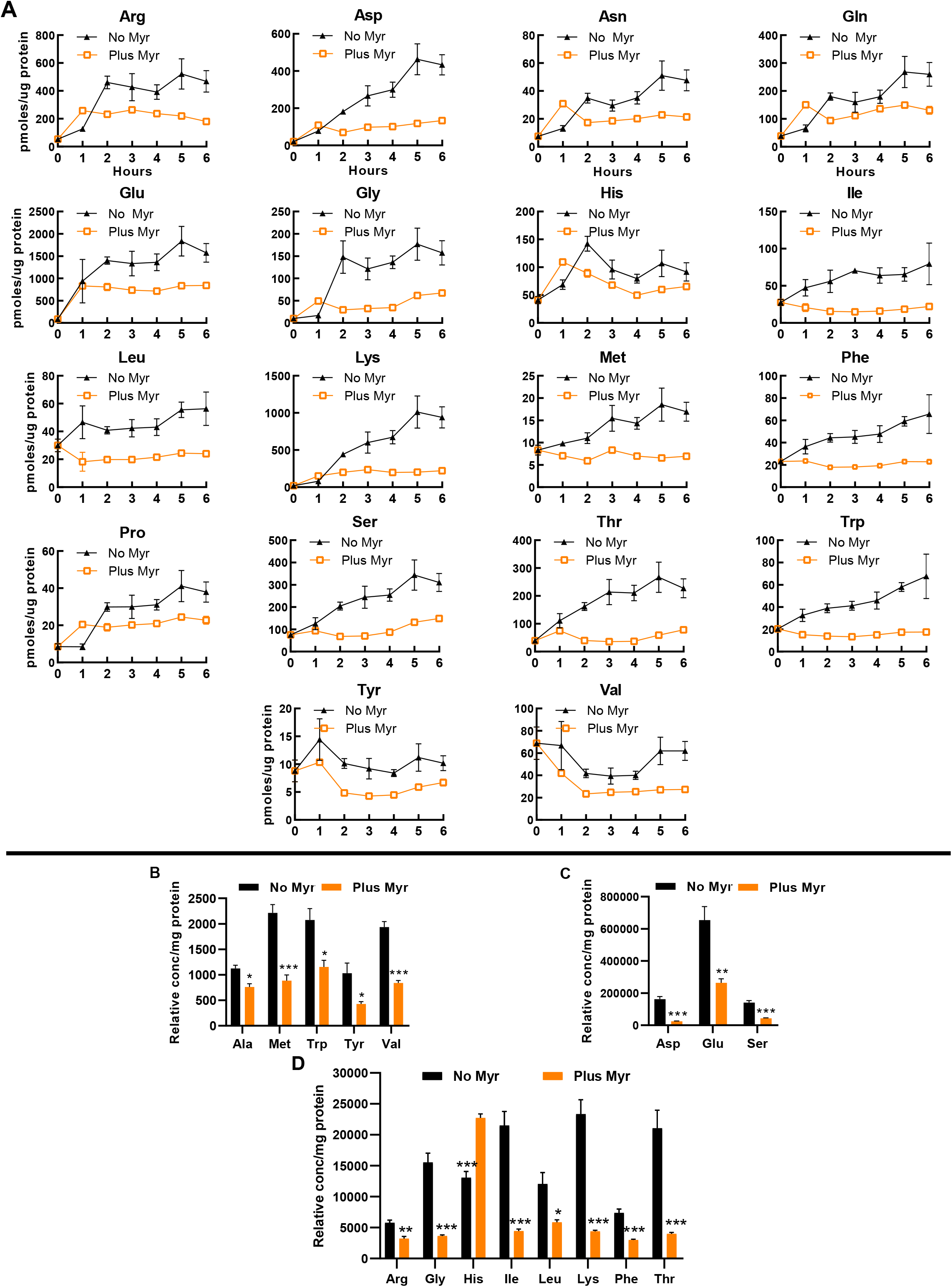
Myr treatment maintains amino acid pools at a low level. (A) Amino acids levels in Myr-treated or untreated cells were measured over a 6 h time span following inoculation of stationary phase cells into fresh SDC medium. Metabolites were analyzed by using the heat-extraction, fluorescent-derivatization procedure. All data were statistically significance by AUC (95% CI: values for each pair of curves do not overlap; error bars are SEM). (B-D) Amino acid levels analyzed by using the GC-MS procedure. Amino acids are grouped into three panels according to their relative concentration. Significance determined by the Student’s *t-test* and error bars are SEM.

To verify free pool data, we quantified amino acids by using gas chromatography-mass spectrometry (GC-MS) as this technology is well suited for examining amino acids [49]. We focus on the 5 h time point because by this time the pool size difference between drug-treated and untreated cells is maximal for most amino acids that can be analyzed by this technique (Asn, Cys, Gln and Pro could not be accurately measured). Samples were analyzed using full-scan mode of the mass spectrometer and ion peaks were normalized to norvaline, added as an internal standard. We present data normalized to the internal procedural standard in order to again emphasize large differences in the size of some amino acid pools (Figure 2B). In both assay procedures, the pools in drug-treated cells are lower than in untreated cells with the exception of His, which is higher at the 5 h time point (Figure 2B) in the GC-MS assay. This difference may result from more effective extraction with the GC-MS procedure. It is also noteworthy that Ala was significantly lower in Myr-treated cells in the GC-MS assay by the 5 h time point (Figure 2B) while there was no statistically significance difference over the 0-6 h assay as measured by AUC.

Lastly, we examined amino acid pools in auxotrophic cells (required uracil, tryptophan, leucine and histidine) grown in strongly buffered SDC medium to eliminate the possibility that weakly buffered medium was somehow necessary for pool lowering in Myr-treated cells. In this culture medium, 14 pools were significantly lowered by Myr treatment (Supplementary Table 1).

Taken together, our results indicate that Myr lowers amino acid pools and extends longevity independent of media acidification.

### Myriocin lowers the rate of amino acid uptake

Since Myr inhibits the production of sphingolipids and thus causes a dramatic change in lipid content at the PM, we hypothesized that the effects of Myr treatment on free pools of amino acids may relate to the function or trafficking of amino acid transporters. Specifically, Myr could reduce amino acid uptake by directly interfering with transport activity, promoting endocytic clearance of amino acid transporters, or impairing their delivery to the PM. To understand the effect of Myr on amino acid transport, we first examined amino acid uptake into cells by using a mixture of ^14^C-labeled amino acids, consisting primarily of glutamate (16%), branched-chain amino acids (29%), alanine (13%) and aspartate (12.5%) with smaller fractions of the other amino acids. Uptake of radiolabel by cells was measured at 4 h when the pool size of most amino acids is significantly lower in Myr-treated cells (Figure 2). To avoid starving cells for specific amino acids in these assays, we quickly harvested cells directly from SDC medium and suspended them in the assay solution containing only the amino acid whose uptake was measured. The concentration of amino acids in assays using the ^14^C-mixture was equal to 11% of the concentration in SDC culture medium to give concentrations slightly below the Km for most AATs [50]. The uptake rate of the ^14^C-amino acid mixture was 52% lower in Myr-treated cells than in untreated cells (Figure 3A), indicating that Myr treatment reduces uptake of several amino acids.

**Figure 3.**
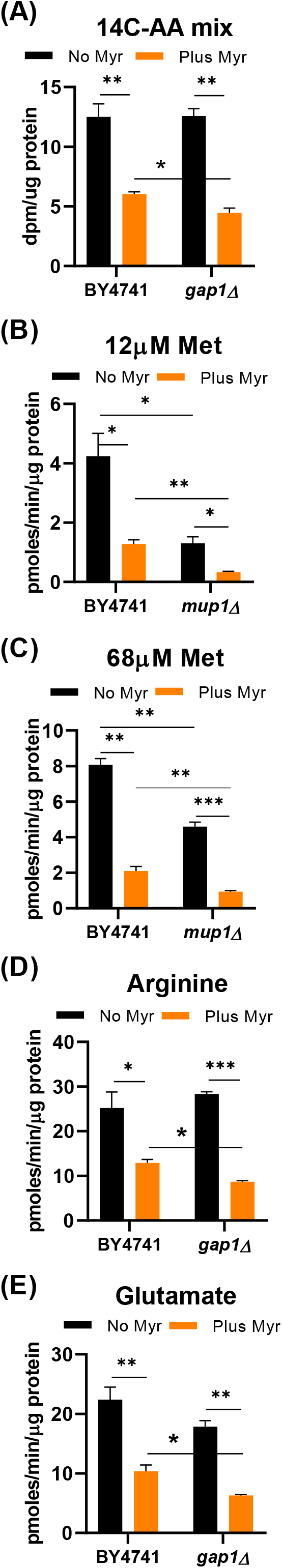
Myr treatment lowers the rate of amino acid uptake. Uptake of radioactive amino acids was quantified after 4 h of growth of the indicated prototrophic strains (BY4741 background) in SDC medium. (A) The concentration of amino acids in the uptake assay was equal to one-fifth of the concentration in SDC medium. The mixture of ^14^C-labeled amino acids contained Glu (16%), branched chain amino acids (29%), Ala (13%), Asp (12.5%) and Phe (8.9%) with smaller amounts of the other amino acids. (B) Uptake of ^3^H-Met was performed with 12 μmol/L substrate, the Km for Mup1 transport, to evaluate activity of this high affinity transporter. (C) Uptake of ^3^H-Met was performed with 68 μmol/L substrate to determine if the activity of other AAT s besides Mup1, with a lower affinity for Met, responded to drug treatment. (D and E). Uptake assays for ^14^C-glutamate and ^14^C-arginine contained 100 μmol/L substrate. Statistical significance determined by using the Student’s *t-test* for all data shown in this Figure and error bars are SD.

The general amino acid permease, Gap1, transports all 20 amino acids and could be responsible for reduced amino acid uptake [51]. To test this hypothesis, we measured uptake of the ^14^C-amino acid mixture in *gap1*Δ cells and found the rate to be identical to that in BY4741 cells without Myr treatment (Figure 3A). However, Myr treatment reduced uptake slightly more in *gap1*Δ than in WT cells (Figure 3A, compare BY4741 to *gap1*Δ). Although the magnitude of difference was slight, the larger reduction of amino acid uptake in Myr-treated *gap1*Δ cells suggests that Gap1 activity contributes to the intracellular pool of amino acids even in the presence of Myr.

Uptake of individual amino acids was examined to identify drug effects on specific AATs. Met uptake was examined because Myr treatment significantly reduces its pool size (Figure 2) and it plays a central role in metabolism and longevity [33, 52–55]. Generally, AATs like the high affinity Met transporter, Mup1, are not highly expressed when substrate is present in culture medium x[56, 57]. Since the SDC culture medium used for most experiments in this study contains a relatively high dose of Met (537 μmol/L, 80 mg/ml), Mup1 expression (and thereby activity) is likely to be low. However, when we compared ^3^H-Met uptake in WT and *mup1*Δ cells using assays containing 12 μmol/L Met, the substrate Km for this transporter [58], Met uptake was relatively robust in WT cells but was reduced about 70% in *mup1*Δ cells (Figure 3B). Thus, in the presence of 12 μmol/L Met, Mup1 is the major Met transporter in BY4741 cells grown in SDC medium. Similar to WT cells, Myr-treatment reduces Met uptake in *mup1*Δ cells to about the same degree, further supporting the idea of other Met AATs are expressed in BY4741 cells and their activity is also reduced by drug treatment.

Met can be transported by six additional AATs [56], with all of them having a higher Km for Met than Mup1 [50]. By increasing the Met concentration in the uptake assay from 12 to 68 μmol/L, we assessed if activity of other Met AATs is lowered by Myr treatment. With 68 μmol/L Met and no drug treatment, the rate of uptake in *mup1*Δ cells is about 40% lower than in BY4741 cells (Figure 3C), supporting the idea that other Met AATs contribute to uptake in conditions of greater Met availability. Met uptake in drug-treated *mup1*Δ cells is reduced by about 55% compared to untreated cells (Figure 3C), further supporting the hypothesis of additional drug-impaired Met AATs being expressed in our experimental conditions. These results indicate that Myr is antagonizing Met uptake by Mup1 as well as other Met transporters.

Several nutrient transporters including Mup1, Can1 (arginine transporter), Lyp1 (lysine transporter, Tat2 (tryptophan and tyrosine transporter) and Fur4 (uracil transporter) are partitioned into a PM domain called the MicroCompartment of Can1 (MCC). MCCs were first associated with endocytosis of PM proteins [59, 60] and now have additional functions [57]. Some of these functions involve sphingolipids and the target of rapamycin complex 2 (TORC2), which is known to play roles in controlling de novo sphingolipid biosynthesis [61–64]. Because of these associations and the low pool of Arg in Myr-treated cells (Figure 2), we examined Arg uptake and found that Myr treatment inhibited uptake of ^14^C-Arg by about 50% (Figure 3D). Only Can1 and Gap1 are known to transport Arg. To determine if one or both of these AATs are present and responding to drug treatment, we examined Arg uptake in untreated *gap1*Δ cells. Uptake is not statistically different in untreated *gap1*Δ and BY4741 cells (Figure 3D), indicating that Can1 is the major Arg transporter under our assay conditions. Myr treatment of *gap1*Δ cells reduces ^14^C-Arg uptake by about 65%. In drug-treated cells there is also a statistically significant lower rate of Arg uptake in *gap1*Δ cells compared to BY4741cells (Figure 3D). From these data we conclude that reduced uptake in both BY4741 and *gap1*Δ cells is due to effects of Myr on Can1 uptake activity.

Lastly, since Myr lowers the free pool of glutamate (Glu) (Figure 2), we examined its uptake, which is only known to occur by the activity of Dip5 and Gap1 [65]. Dip5 is a dicarboxylic acid transporter that also transports aspartate, Gln, Asn, Ser, Ala, and Gly [50, 65], all of which exhibit decreased levels in cells treated with Myr. Notably, Dip5 is not present in MCC domains [57] and thus, provides insights into whether the effects of Myr are restricted to MCC-localized AATs. Uptake of ^14^C-Glu in BY4741 cells is lowered about 50% by Myr-treatment (Figure 3E). To examine the effect of Myr treatment on Dip5-mediated Glu transport, we measured ^14^C-Glu uptake in *gap1*Δ cells and found a 65% decrease in uptake rate (Figure 3E). These data indicate that a majority of Glu is transported by Dip5 and its activity is lowered by Myr treatment. Importantly, these data indicate that the effects of Myr are not limited to AATs associated with MCCc at the PM.

### Myriocin reduces Mup1 transport activity in the plasma membrane

Trafficking of Mup1 from its site of synthesis at the endoplasmic reticulum through the secretory pathway to the PM and eventual endocytosis by a ubiquitin-mediated process have been studied extensively [57, 66]. These previous studies provide valuable insights into the regulation of Mup1 trafficking and transport activity in the context of changing amino acid or nitrogen availability. We set out to understand how Myr reduces Mup1 activity and to determine if this affect is caused by ***(i)*** reduced delivery of Mup1 to the PM via the secretory pathway, ***(ii)*** increased removal of Mup1 from the PM by endocytosis, or ***(iii)*** direct inhibition of Mup1 transport activity at the PM.

To quantitatively evaluate the effect of Myr treatment on Mup1 protein level and cellular location, we examined cells with the chromosomal *MUP1* gene fused to the green fluorescent protein (GFP) to determine if drug treatment impaired trafficking of Mup1-GFP to the PM. Starting with log phase cells treated with Met to deplete Mup1-GFP from the PM (Figure 4A, top-left panel, +Met) cells were shifted to methionine-deficient SC medium to promote synthesis of Mup1-GFP in the absence or presence of a range of Myr concentrations. Trafficking of Mup1-GFP to the PM 1 h after the media shift was not impaired by any concentration of Myr (Figure 4A). In these assays Vph1-MARS served as a visual marker for the vacuolar membrane to track the internal pool of Mup1-GFP localized to the lumen of the vacuole. Consistent with these observations, we also found that yeast cells grown to stationary phase (which triggers Mup1 clearance from the PM) [67] and inoculated into methionine-deficient SC medium exhibited delivery of newly-synthesized Mup1-GFP to the PM within 1 h, which was unaffected by Myr treatment (Figure 4B). From these data we conclude that Myr does not impair trafficking of newly synthesized Mup1 to the PM via the secretory pathway.

**Figure 4.**
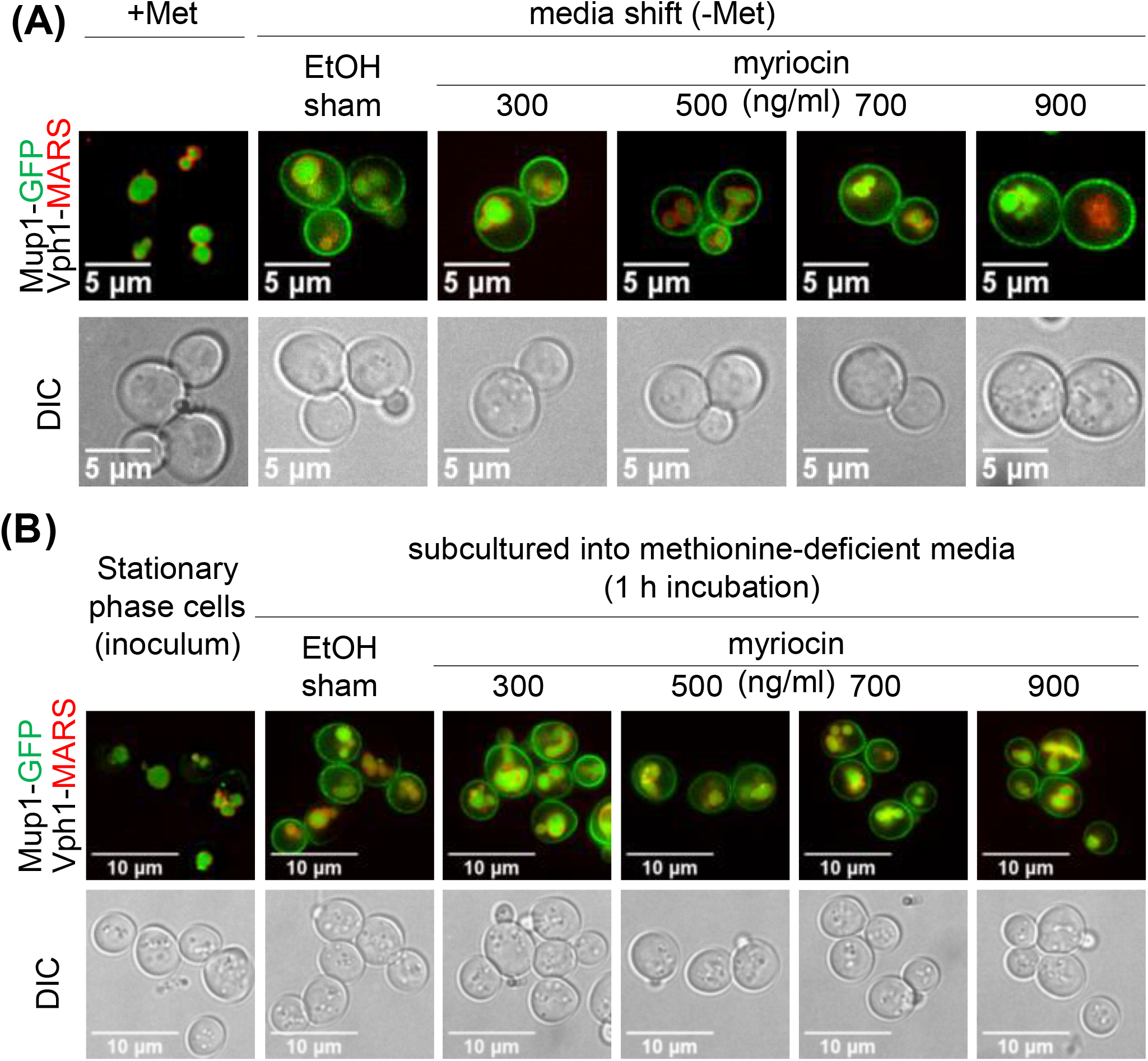
Mup1 biosynthesis and secretion appear unaffected by myriocin treatment. (A) JTY240 cells, with chromosomal copies of *MUP1-GFP* and *VPH1-MARS,* were gown to mid-log phase (A600nm = 0.4) and treated with Met (0.134 umol/L or 20 μg/ml) for 1 hour to deplete Mup1-GFP at the PM. Cells were washed with water, shifted to Met-deficient SC medium with or without Myr treatment for 1 hour and visualized by fluorescence microscopy. (B) JTY240 cells grown overnight in SC medium to stationary phase were washed with water and cultured (starting A600nm = 0.15) in fresh SC medium with or without Myr treatment.

The media-shift experiments presented in Figure 4 allow for sensitive detection of Mup1-GFP delivery to the PM, but because of the media and culturing protocol cells exhibit a high vacuolar GFP signal at the beginning of each experiment, making it difficult to determine if Myr induces the endocytosis of Mup1-GFP. To assess Mup1 endocytosis, we examined cells harboring chromosomal *MUP1* tagged with ecliptic pHluorin (a GFP variant that loses fluorescence at low pH) which does not fluoresce in the vacuole and late endosomal compartments and thus can be used to quantify the abundance of Mup1 at the PM and early endosomal compartments [68–70]. Stationary phase cells inoculated into fresh, methionine-deficient SC medium exhibited a steady increase in Mup1-pHluourin signal over a 9 h culture time course (Figure 5A-B), consistent with results observed in Figure 4. However, inoculation into Myr-containing media resulted in an initial increase in Mup1-pHluorin at 3 h followed by a sharp decline in Mup1-pHLuorin by 6 h (Figure 5A-B). This Myr-induced decline in Mup1-pHlourin signal at later time points was also detected in flow cytometry analysis (n≥10,000 cells per time point). Specifically, following transfer of stationary phase cells into fresh methionine-deficient medium, we observed very a low Mup1-pHluorin signal that increased until plateauing at about 4 h (Figure 5C). This early phase of Mup1-pHluorin synthesis and delivery to the PM was unaffected by Myr treatment (Figure 5C), consistent with the results of Figure 4. After plateauing around the 4 h time point, the Mup1-pHluorin signal began to decrease slowly (Figure 5C), suggesting that Mup1-pHluorin PM abundance had reached a steady state balanced by endocytic clearance. Importantly, the Mup1-pHluorin signal decreased more rapidly in the presence of Myr, an effect that was dose-dependent (Figure 5C).

**Figure 5.**
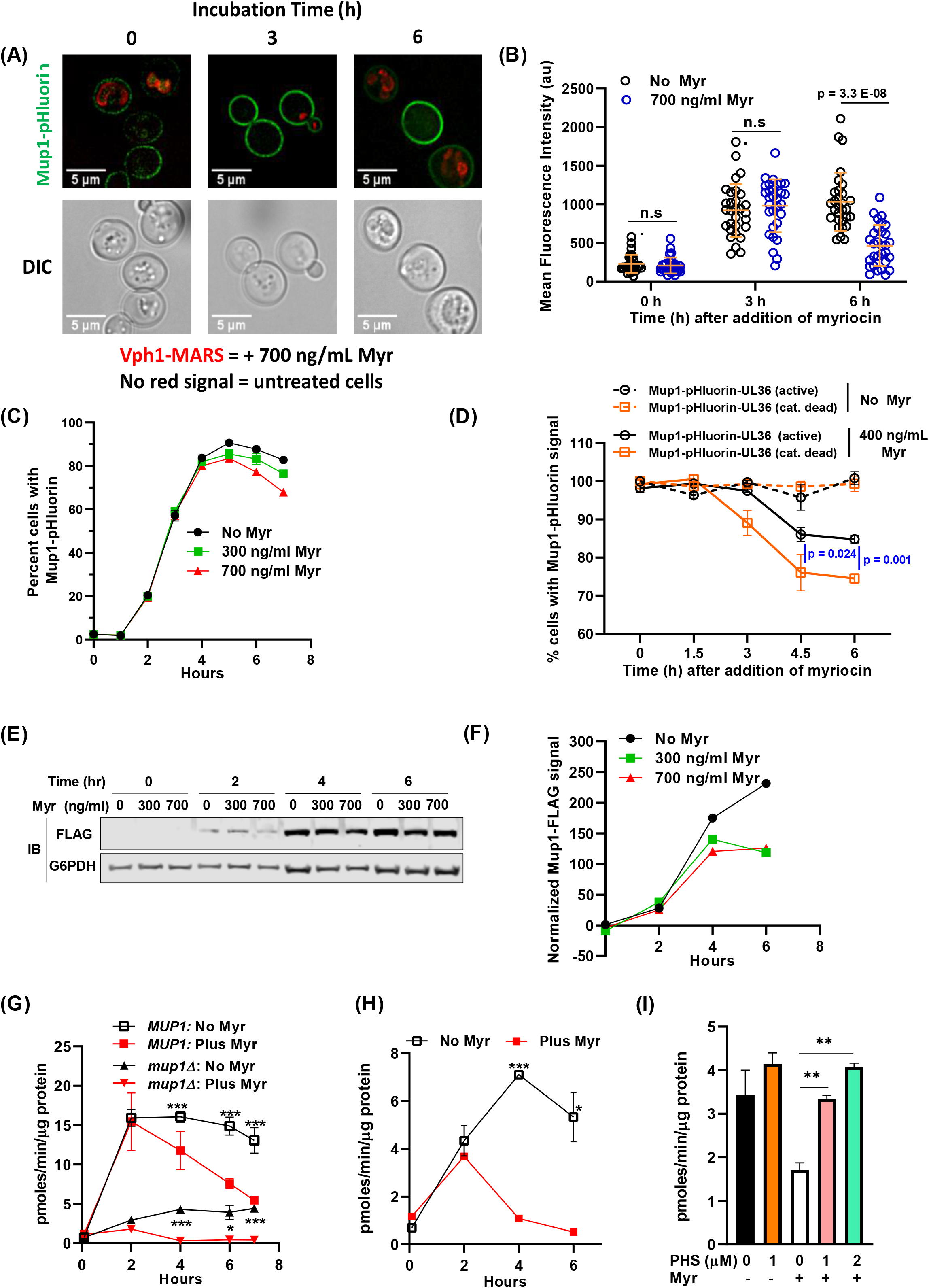
Myr treatment reduces Mup1 activity in the PM. (A) Cells expressing Mup1-pHluorin (NHY413) or Mup1-pHluorin and Vph1-MARS (NHY414) grown for 20 h to stationary phase and then inoculated into methionine-deficient media in the presence (NHY414) or absence (NHY413) of 1.743 umol/L (700 ng/ml) Myr. Cells were mixed prior to visualization by fluorescence microscopy so that untreated and Myr-treated cells can be visualized side-by-side. (B) Quantification of Mup1-pHluorin microscopy signal shown in (A). Mean signal intensity of Mup1-pHluorin at the plasma membrane was measured for individual cells (n≥30) using ImageJ-Fiji. (C) Relative fluorescence intensity of Mup1-pHluorin in JMY1811 cells was determined over time in triplicate cultures by flow cytometry. Cells grown to stationary phase in SC medium lacking Met were washed, diluted into fresh medium, and 10,000 cells were analyzed at each sampling time point in cultures containing or lacking Myr. (D) Cells expressing Mup1-pHluorin-UL36 (NHY447) or Mup1-pHluorin-UL36 catalytic dead Vph1-MARS (NHY431) were grown to mid-log phase, mixed, and subcultured into methionine-deficient media with or without 0.996 μmol/L (400 ng/ml) Myr. Fluorescent cells were quantified in 5000 sampled cells over time using a flow cytometer. (E) Immunoblotting analysis of Mup1-FLAG in cells grown in SDC medium. G6PDH immunoblots (bottom panel) were used as a loading control for the blots. (F) Quantification of the anti-FLAG immunoblot signal shown in (E) using the relative fluorescence of G6PDH as a loading normalization. (G) The uptake rate of 12 μmol/L ^3^H-Met in JMY1811 and JMY305 *(mup1*Δ) cells was measured on cells grown in the same way as described in panel (C). Statistical significance *MUP1:No* Myr vs Plus Myr (AUC, 95% CI: 88.03-97.24 vs 56.89-80.86) and *mup1:No* Myr vs Plus Myr (AUC, 95% CI: 21.02-25.09 vs 5.593-6.345). (H) The uptake rate of 12 μmol/L ^3^H-Met was measured on JMY1811 cells grown in the same way as described in panel (E). Statistical significance: No Myr vs Plus Myr (AUC, 95% CI: 26.02-31.38 vs 10.67-11.38). (I) Supplementing culture medium with phytosphingosine (PHS) restores Met uptake in Myr-treated BY4741 cells. ^3^H-Met uptake was measured with a substrate concentration of 12 μmol/L in cells grown for 4 h in SDC medium with or without PHS and with or without Myr treatment.

To test if the loss of Mup1-pHluorin signal following Myr treatment is due to ubiquitin-mediated endocytosis, we generated yeast cells with chromosomal *MUP1* tagged with pHluorin-UL36 (catalytic active and catalytic dead). UL36 is a viral deubiquitylase enzyme (DUB) that can reverse localized ubiquitin conjugation activity, thus protecting the fusion protein from ubiquitin-mediated degradation or trafficking events [71]. Importantly, Myr-treatment induced the clearance of a catalytic dead Mup1-pHluorin-UL36 while the catalytic active variant was partially stabilized (Figure 5D). These results indicate that ubiquitin-mediated endocytosis is at least partially responsible for the loss of Mup1-pHluorin signal following Myr treatment, although this does not exclude the possibility that other clearance mechanisms may contribute to the response. Taken together, these observations indicate that Myr does not affect Mup1 biosynthesis or delivery to the PM but can promote its endocytic clearance from the PM following prolonged exposures.

We next sought to analyze Mup1 trafficking in cells grown in SDC medium containing methionine (537 μmol/L). As expected, the Mup1-pHluorin signal is very low in cells grown in SDC medium and potential effects of Myr treatment are not observable (Figure S5). To further characterize the effects of Myr treatment we examined Mup1 protein abundance in cells expressing Mup1-3xFLAG (tagged at the chromosomal locus) by immunodetection of Mup1-3xFLAG in cell lysates (Figure 5E-F). There is no detectable Mup1-FLAG at time 0 when stationary phase cells are inoculated into fresh SDC medium (Figure 5E, left lanes, top panel). After growth for 2 h similar FLAG-signals were detected in Myr-treated or untreated cells (Figure 5E). However, while Mup1-FLAG steady state abundance continues to increase through 6 h in untreated cells, its abundance plateaus at 4 h in Myr-treated cells, consistent with increased endocytic clearance at later time points in this experiment (Figure 5E-F). Thus, even in culture medium containing a fairly high concentration Met, the effects of Myr treatment on Mup1 abundance are similar to those observed in methionine-deficient SC medium (compare data in Figures 5C and 5E-F).

These observations, coupled with the observed reduction in ^3^H-Met uptake (Figure 3), prompted us to correlate the kinetics of methionine transport with Mup1 trafficking over the course of a 7-hour Myr treatment period (as shown in Figure 5C). The rate of Met uptake is very low following inoculation from a stationary phase culture (Figure 5G), which was expected since Mup1 is not expressed in cells at stationary phase. However, Met uptake capacity increased rapidly over 2 h of growth and to the same degree in cells with and without Myr treatment (Figure 5G). After 2 h of growth, Met uptake capacity slowly decreased in untreated cells, while uptake capacity decreased to a much greater extent in Myr-treated cells. Importantly, the majority of this uptake activity can be attributed to Mup1, since uptake of ^3^H-Met was reduced by ~70% in *mup1*Δ cells (Figure 5G). Interestingly, Myr treatment reduced ^3^H-Met uptake even in *mup1*Δ cells (Figure 5G), consistent with data shown in Figure 3 indicating that other transporters also contribute to Met uptake in these experiments.

We hypothesized that decreased Met uptake during Myr treatment may be due to Myr-induced endocytic clearance of Mup1, however such clearance was only observed in later time points following Myr treatment. Alternatively, it is also possible that Myr decreases Met uptake by directly inhibiting Mup1 transport activity at the PM. To distinguish between these possibilities, we examined ^3^H-Met uptake activity of cells grown in SDC medium and observed, as in cells grown in SC medium, very low uptake at time 0 in cells with and without Myr treatment (Figure 5H). In untreated cells, ^3^H-Met uptake increases steadily during 4 h of growth in SDC, followed by a slight decrease in Met uptake observed at 6 h (Figure 5H). In contrast, Myr-treated cells exhibit an initial increase in Met uptake at 2 h of growth followed by a significant decline in Met uptake to levels observed at the zero-time point (Figure 5H) despite the observed increase in Mup1 abundance during that time frame (Figure 5A-C and 5E-F). This disparity between increased Mup1 abundance and decreased Met uptake in Myr-treated cells indicates that Myr treatment reduces Mup1 transport activity prior to stimulating endocytic clearance from the PM.

Based on these data, we hypothesized that Myr treatment gradually reduces the concentration of sphingolipids in membranes, which reduces Mup1 transport activity. To test this hypothesis, we analyzed Met uptake in culture medium supplemented with phytosphingosine, a downstream product of serine palmitoyltransferase (Figure 1A, SPT). Such supplementation is an established way to bypass both pharmacological and genetic impairment of SPT activity [72, 73]. While phytosphingosine had no effect on ^3^H-Met uptake in cells that were not treated with Myr, it restored uptake in Myr-treated cells (Figure 5I). Taken together, these results indicate that Myr-treatment induces a loss of Mup1 transport activity followed by endocytic clearance from the plasma membrane.

### Sensing of amino acid pool size by TORC1 is vital for increased lifespan

An evolutionarily conserved function of TORC1 is to adjust growth and proliferation to a rate commensurate with amino acid levels [38, 43, 74]. For this reason, amino acid abundance promotes TORC1 activity while amino acid scarcity reduces it. Accordingly, impairment of amino acid pool expansion by Myr treatment (Figure 2) is likely a key upstream regulatory mechanism for slowing the growth rate after 4 h of drug treatment (Figure 1E) and for reducing TORC1 activity as we noted previously [39]. Because reduced TORC1 activity is a prominent and conserved feature of longevity [25, 26, 38], we determined if lower amino acid pool size in drug-treated cells is sensed by TORC1 to reduce its activity and promote CLS.

The Gtr1/Gtr2 GTPase signaling pathway in yeast (RagA or RagB/RagC or RagD in mammals) senses amino acids and regulates TORC1 activity [75–77]. Hence, amino acids promote GTP loading onto Gtr1/2 which activates TORC1 [78]. Conversely, amino acid scarcity turns off Gtr1/2 by causing Iml1, a GTPase-activating protein (GAP) found in a complex with Npr2/Npr3 (SEACIT complex, GATOR1 in mammals), to hydrolyze GTP and inactivate Gtr1/2 which reduces TORC1 activity. We reasoned that deleting *IML1 or NPR2* should prevent TORC1 from sensing the smaller amino acid pools in drug-treated cells because Gtr1/2 would remain active, promote TORC1 activity and impair lifespan enhancement. Reduced Myr concentrations were required for lifespan assays to partially mitigate the slower growth rate of *iml1*Δ and *npr2*Δ cells and reduced final cell density caused by drug treatment. With reduced Myr concentrations, growth rate and final cell density are slightly reduced compared to BY4741 cells (Figure 6A). We find that untreated *iml1*Δ and *npr2*Δ cells lose viability slightly faster than WT BY4741 cells (Figure 6B and 6C). Notably, Myr treatment does not increase the lifespan of *iml1*Δ and *npr2*Δ cells. These data support the hypothesis that the Gtr1/2 pathway is essential for sensing lowered amino acid pools, down-regulating TORC1 activity and increasing lifespan in Myr-treated cells.

**Figure 6.**
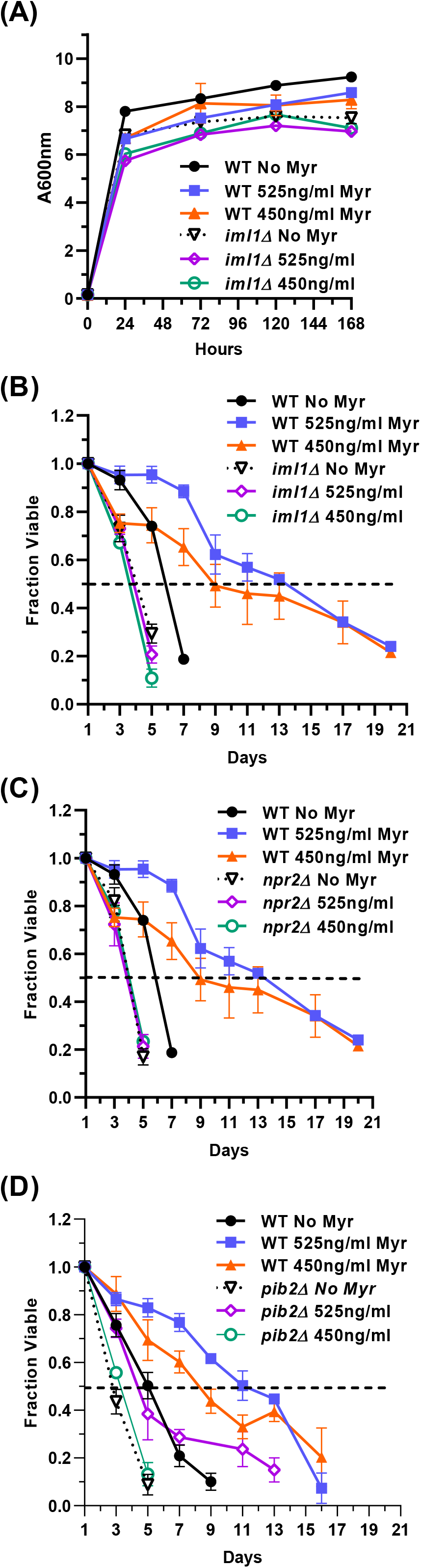
The Pib2 and Gtr1/2 pathways sense amino acid pools to control TORC1 activity and increase lifespan in Myr-treated cells. (A) Growth rate and cell density of indicated strains with or without Myr treatment. (B and C) The role of the Gtr1/2 pathway in lifespan was examined by measuring the viability of prototrophic WT BY4741, *iml1*Δ and *npr2*Δ cells, with and without Myr treatment, over time. (D) The role of the Pib2 pathway in lifespan was examined by measuring the viability of prototrophic BY4741 and *pib2*Δ cells, with and without Myr treatment, over time.

A yeast pathway for sensing glutamine and leucine upstream of TORC1 includes Pib2, which harbors a PI3P-binding FYVE-domain, and Vps34 which synthesizes this lipid signal [79–81]. We determined if the Pib2 pathway plays a role in Myr-enhanced lifespan by using *pib2*Δ cells which are not as sensitive to Myr treatment as *iml1*Δ and *npr2*Δ cells. In a lifespan assay, untreated *pib2*Δ cells die slightly faster than WT BY4741 just like *iml1*Δ and *npr2*Δ cells (Figure 6D). However, there is a small difference because *pib2*Δ cells tend to respond to Myr since the 1.309 uM (525 ng/ml) dose slightly increases survival by a small but statistically significant amount relative to *pib2*Δ cells without drug treatment (AUC: 95% CI, 1.672-2.247 vs 2.980-4.104). These results are consistent with data showing that the Vps34-Pib2 and Gtr1/2 pathways cooperate to control TORC1 activity [82, 83].

## Discussion

Aging is an established risk factor for many chronic diseases, several contributing heavily to mortality in the elderly [13]. Available data indicate ways to modulate aging and improve health in later life. The best characterized approaches seek to manipulate nutrient intake in ways that increase lifespan in model organisms and improve health metrics in humans [4]. Regrettably, most humans are unlikely to achieve health benefits because these approaches require long-term restraint of our inborn desire to eat.

The work described here contributes to the field of longevity research by establishing a proof-of-principle that a pharmacological agent can increase lifespan by inducing a physiological state resembling amino acid restriction that modulates the conserved TORC1 pathway to promote longevity. Worms, flies and mammals including humans are very likely to have been exposed to food contaminated with myriocin-like compounds, particularly their gut epithelial cells, which could have evolved a mechanism to lower the free pool of at least some amino acids so as to induce cytoprotection and stress responses and increase survival [84, 85]. Such myriocin-induced, hormesis-like stress responses have been demonstrated in *Caenorhabditis elegans* [86] and recently Myr treatment, along with RNAi knockdown or mutation of the serine palmitoyltransferase gene, were shown to increase lifespan in *C. elegans* [87]. Finally, a growing body of data show that Myr treatment reduces the severity of age-related diseases in mice and rats: low dose Myr treatment lowers atherosclerosis and cardiac impairment [88–91] and reduces risk factors for metabolic syndrome, obesity, diabetes and cancer [92–96]. Myr also reduces amyloid beta and tau hyperphosphorylation in a mouse model of Alzheimer’s disease [97] and has therapeutic effects on other neurodegenerative diseases [98, 99]. These results from multi-cellular eukaryotes combined with the data presented here strongly support the notion that Myr-like compounds could mitigate debilitating effects of age-related diseases in humans. Thus, it will be critical to understand the mechanistic basis of Myr-induced longevity.

Here, we report that Myr induces cells to enter a quiescent or Q state (Figures 1C and 1D) which is known to enhance longevity [43, 100, 101]. Budding yeast have been an excellent model for understanding quiescence. Starvation for assorted nutrients, while taking unique routes to promote the quiescent state, have common features including increased stores of glycogen, trehalose and triglycerides, down-regulate the TORC1 and PKA pathways and increased AMPK/Snf1 signaling along with autophagy [43, 102–104]. Significantly, we previously found Myr-treated cells possess all of these features [14, 39]. Notably, down-regulation of TORC1 is also necessary for mammalian cells to enter a quiescent state [45].

The ability of Myr to maintain at least 17 amino acid pools (exceptions: Ala and His; Cys not analyzed) at a low level (Figure 2) is unexpected and unique. Analysis of amino acid pools in the yeast gene-deletion strain collection did not reveal any strain with as many reduced amino acid pools [40] as we find with Myr treatment. This comparison suggests that Myr treatment influences multiple genes and signaling pathways. Data for amino acid uptake (Figure 3) support drug impairment of several AATs by Myr, including Met uptake by Mup1 and by one or more of the other transporters capable of transporting Met such as Mup3, Agp3, Agp1, Bap2, Bap3 and Gnp1 [56]. In addition, uptake assays with ^14^C-arginine and ^14^C-glutamate, implicate Can1 and Dip5, respectively (Figure 3D and 3E), as additional AATs whose activity is impaired by Myr treatment. Lastly, assays using a mixture of ^14^C-labeled amino acids and comparing uptake in BY4741 with *gap1*Δ cells, show that Gap1 does not transport the majority of amino acids in this mixture, thus, transport requires other AATs including ones transporting branched chain amino acids (Leu, Val and Ile) along with Asp, which make up 41% of the amino acid mixture and whose pools are low in drug-treated cells (Figure 2).

Loss of Mup1 transport activity in Myr-treated cells may be related to the reported loss of Gap1 transport activity in cells with a temperature-sensitive *lcb1-100* gene. This mutant allele slows the rate of sphingolipid synthesis at restrictive temperatures [73]. A lack of transport activity is a key feature of Gap1 synthesized after cells are shifted to a restrictive temperature. The authors hypothesized that the lack of transport activity was due to reduced sphingolipid synthesis caused by reduced enzymatic activity of the temperature-sensitive serine palmitoyltransferase (the target of Myr) present in *lcb1-100* cells. Support for this hypothesis rested on data showing restoration of Gap1 activity in *lcb1-100* cells treated with a long-chain base, the product of serine palmitoyltransferase (Figure 1A, SPT), which enabled cells to make sphingolipids even at a restrictive temperature. In accordance with these studies, we find that supplementation of culture medium with a long-chain base (phytosphingosine) bypasses the effect of Myr treatment on ^3^H-Met transport restoring uptake to the level observed in untreated cells (Figure 5I). Such restoration of Mup1 activity is strikingly similar to the restoration of Gap1 activity in *lcb1-100* cells. Since we found evidence for Myr-impairment of Can1, Dip5 and other Met transporters besides Mup1, it is tempting to speculate that the transport activity of many AATs, perhaps most, are likewise impaired by drug treatment as this would explain the widespread reduction in pool size we observe (Figure 2).

TORC1 relies on several pathways to sense amino acids and the two that we examined, Gtr1/Gtr2 and Pib2, appear to be necessary for cells to respond to Myr treatment and promote longevity (Figure 6). Pib2 is known to sense glutamine and leucine [83], consistent with their reduction by Myr treatment (Figure 2). Gtr1/Gtr2 sense several amino acids and the specific one or ones promoting longevity in Myr-treated cells will require further study.

Additional amino acid sensors that could be controlling TORC1 activity [105] in Myr-treated cells need to be examined in the future as some could sense specific amino acids and modulate key metabolic and physiological functions necessary for Myr-induced longevity. One such key amino acid is Met with its established roles in longevity [28–33]. Limiting Met availability could reconfigure cellular physiology in many ways to enhance longevity. For instance, Met limitation during overall amino acid lowering reprograms anabolic metabolism to match nutritional status with proliferation [106]. Additionally, a lowered Met level was shown to be a key control point for replicative lifespan, with and without glucose restriction, in yeast [33]. Cross-talk between glucose restriction and Met metabolism was proposed to be a general mechanism to balance nutrient availability with translation and growth. Thus, it is conceivable that the lowered Met pool in Myr-treated cells plays a central role in lifespan enhancement.

Based on data presented in this manuscript, we hypothesize that Myr treatment decreases amino acid pools by two mechanisms: (1) an acute decrease in the function of amino acid transporters, caused by decreased sphingolipid concentrations in membranes, and (2) increased endocytic clearance and vacuolar degradation of amino acid transporters following prolonged exposure to Myr. This latter effect may be related to misfolding of AATs, caused by changes in lipid composition at the PM and sensed by PM protein quality surveillance machinery [107]. Alternatively, the induction of endocytic clearance at later time points following exposure to Myr (Figure 5B and 5C) may suggest that this occurs as part of a cellular adaptive response, perhaps related to amino acid starvation. Additional experimentation will be required to distinguish between these possibilities.

Like rapamycin, myriocin was initially studied for its immunosuppressive activity produced at high dosages [19]. And like rapamycin, lower doses of myriocin have been shown to ameliorate age-related diseases [90, 91, 98, 108]. While humans might not tolerate long-term use of myriocin-like compounds, understanding how they enhance physiological functions and lifespan in model organisms by lowering multiple amino acid pools, could uncover new avenues for drug development. Unlike restriction for a single amino acid, lowering multiple amino acid pools may have effects more related to total protein restriction, which enhances lifespan in organisms ranging from yeasts to mammals [6, 7, 109–111]. Furthermore, recent studies of caloric restriction implicate reduced intracellular methionine in yeasts [33] and glycine-serine-threonine metabolism (folate and methionine cycles and the transulfuration pathway) in mice [112] as key drivers of longevity. Thus, it’s reasonable to propose that the reduction of amino acids in one-carbon and related metabolism (Gly, Ser, Met and Thr) that we observe in myriocin-treated cells, are mediating at least some of the longevity increase. Given the complexity of aging and of cellular metabolism, reduction of other amino acids by myriocin including the branched-chain amino acids Leu, Ile and Val may contribute to longevity. Future studies will aim to identify the primary mediators of myriocin-induced longevity which may include other mechanisms in addition to lowered amino acid pools [39].

## Materials and methods

### Strains, culture conditions, lifespan assays and statistical significance

Yeast strains are described in the Table 1. BY4741 cells and mutant strains from the BY4741 gene knockout library were made prototrophic by transformation with the pHLUM plasmid (Addgene, Watertown, Massachusetts) [40]. SDC culture medium and procedures for CLS assays followed published procedures [117] except for the following changes: culture medium was warmed to 30°C in 125 ml or 1 L flasks. The order of additions was: 95% EtOH, Myr (to give a final concentration of 0.3% EtOH) and then cells to give an initial A600nm units/ml of 0.15 or as indicated in the text. After each addition, cultures were mixed for 10-15 sec. Cultures were incubated in a gyratory shaker (200 rpm) at 30°C for 72 h (day 1 of CLS assay) and viability was quantified by diluting cells in sterile water, plating on YPD plates and counting colonies after 3 days incubation at 30°C. Colony values were plotted as ‘Fraction Viable’ with day 1 set to a value of 1, and subsequent days as a fraction of the day 1 colony count. Myriocin (Cayman Chemicals, #63150) stock solutions of 2.49or 0.996 mM (0.4 mg/ml) in 95% EtOH were treated before use to three cycles of 1 min at 65°C followed by 1 min vortexing. Myr stocks were stored at −20°C. Phytosphingosine (MilliporeSigma, P2795), dissolved in 95% EtOH to give a 20 mM stock, was added to cultures after addition of EtOH and before addition of Myr. The combination of phytosphingosine and Myr slows growth, therefore, we used 1.246 μM (500 ng/ml) of Myr for the uptake assays shown in Figure DI. In experiments conducted using SC media, cells were maintained and cultured as previously described [69].A two-tailed Student’s t-test (*= ρ<0.05; **= ρ<0.01; ***= ρ<0.001) and the Area Under the Curve (AUC, Prism 8, Graphpad) were used to evaluate statistical significance of CLS and other data representing at least 3 biological replicates. Results were verified by at least one repeat experiment.

**Table 1.** Strains used in this study.

| Strain | Genotype | Reference |
| --- | --- | --- |
| BY4741 | *MATa his3-Δ1 leu2-Δ0 ura3-Δ0 met15-Δ0* | [113] |
| *pib2Δ* | BY4741 with *pib2::KAN* | Horizon [114] |
| *npr2Δ* | BY4741 with *npr2::KAN* | Horizon [114] |
| *iml1Δ* | BY4741 with *iml1::KAN* | Horizon [114] |
| SEY6210 | *MATalpha leu2-2,112 ura3-52 his 3delta200 trp1-delta901 lys2-801 suc2-delta9* | [115] |
| JMY1811 | SEY6210 with *Mup1::pHluorin-KanMX* | [116] |
| JTY240 | SEY6210 with *Mup1::GFP-KanMX* and *Vph1::Mars-TRP1* | [70] |
| JMY305 | SEY6210 with *mup1::TRP1* | This study |
| JMY78 | *SET6210 with MUP1::6xHis-TEV-3xFLAG-TRP1* | This study |
| NHY413 | *SEY6210 Mup1-pHluorin::NATMX* | This study |
| NHY414 | *SEY6210 Mup1-pHluorin::NATMX Vph1-MARS::TRP1* | This study |
| NHY431 | *SEY6210 Mup1-pHluorin-UL36(N-term 15-260 HSV1 UL36, active)::KANMX* | This study |
| NHY447 | *SEY6210 Mup1-pHluorin-UL36(N-term 15-260 HSV1 UL36 C40S, inactive)::KANMX* | This study |

To understand how Myr reconfigures cellular physiology to enhance survival requires time course assays and larger numbers of cells than are available from typical 25 ml cultures used for CLS assays. In addition, a higher starting density than 0.005 A600nm units of cells is needed to access early changes induced by Myr treatment (Figure 1B). By varying the starting culture density and the concentration of Myr, we found conditions in which Myr stimulated CLS to approximately the same extend in 200 ml cultures (1 L flask) started at an A600nm of 0.15 (Figure 1G), as in 25 ml cultures started at an A600nm of 0.005 (Compare Figures 1B and G). One limitation of lifespan assays performed on larger cultures and with prototrophic cells is a tendency for mutant cells that feed off of dead cells, to multiply and produce an increase in viability as noted by others [24]. This is what occurred with two of the three drug-treated cultures around day 15 of the experiment shown in Figure 1G and is why no further data points are presented. The amino acid pool and uptake assays and Mup1 protein trafficking data presented here are performed long before re-growth begins and are not affected by it.

### Rationale for drug doses

Myriocin has been used in a wide variety of studies and in organisms ranging from yeasts, worms and flies to mammals [14, 92, 99]. In contrast to some studies which use an acute drug dose to induce on-off types of physiological responses, our studies use a moderate drug dose or chronic treatment to induce relatively small, hormesis-like stress and related physiological responses. To improve reproducibility across different types of assays and strain backgrounds, the activity of drug stocks was quantified from the mass yield [118] measured at A600nm after 24 h of growth in SDC containing. A concentration of drug that reduced the A600nm by 20-25% was used for assays (typically 1.245-1.745 μM (500-700 ng/ml)).

### Metabolite analyses

A protocol for measuring free amino acid pools was modified to enable time-course assays [78]. Flasks (1 L) containing 200 ml of SDC were warmed to 30°C and EtOH +/- Myr was added as described for CLS assays. Starter cultures of prototrophic BY4741 cells, grown for 24 h in SDC at 30°C, were diluted into the 200 ml cultures to an A600nm of 0.15 units/ml. Flasks were incubated at 30°C on a gyratory shaker at 200 rpm. Every hour for 6 h, 5 A600 units of cells were collected on membrane filters (0.45 μm, HAWP02500, MF-Millipore, MilliporeSigma, Burlington Massachusetts). Filters were immediately rinsed twice with 5 ml of ice-cold Nanopure water, quickly transferred to 0.5 ml cold water, vortexed and incubated 15 min in a boiling water bath. Cells were chilled on ice for 5 min and centrifuged 1 min at 13,000 x g. Fifty microliter of supernatant was saved for protein assay (BCA assay, Pierce) while the rest was filtered (Pall Nanosep^®^ OD003C33 centrifugal device with 3K Omega membrane, Pall Corp.) to remove molecules larger than 3,000 Daltons. Filtrates were transferred to screw-capped tubes, frozen in a dry ice/ethanol bath and stored at −80°C. Amino acid analyses were done at the Proteomics and Mass Spectrometry Facility at the Donald Danforth Plant Science Center (St. Louis, Missouri, USA). Data are expressed as pmoles/ug of protein.

For amino acid analyses by GC-MS, cells were grown as in the heat extraction method, but they were collected on nylon membrane filters (Whatman, 0.45 μm, 25 mm dia., WHA7404002) followed by washing once with 5 ml ice-cold Nanopure water. Filters were immediately immersed into 2 ml of icecold acetonitrile:water (1:1, v/v, containing 20 μM L-norvaline as an internal standard) in a 5 ml capped, conical centrifuge tube and vortexed 10 sec before removing the filter. Samples were shaken up and down sixty times and incubated on ice for at least 20 min before freezing in liquid nitrogen and storing overnight at −80°C. Tubes thawed on ice, were vortexed 3 min at room temperature followed by centrifugation at 4,500 x g for 5 min at 4°C (Eppendorf #5010R). The supernatant fluid was poured into a chilled 2 ml screw-capped tube, centrifuged again and equal amounts were transferred to microfuge tubes and dried at 10^-3^ mBar using a Labconco CentriVap system (ThermoFisher Scientific). Dried samples were either immediately derivatized and analyzed by GC-MS or stored at −80°C. Cells pellets washed once with 1 ml cold methanol (−20°C) and air-dried. Cell pellets were suspended in 250 μl of 2% sodium dodecyl sulfate and 5 mM Tris-HCl (pH 7.4) overnight at 4°C. Following addition of 250 μl of water, the samples were stored at −20°C. Thawed cell pellets were heated 5 min at 95°C before measuring protein contend by using a micro-BCA assay (Pierce, ThermoFisher Scientific) and a NanoDrop One spectrophometer (ThermoFisher Scientific).

### Sample derivatization

Extracted samples were derivatized by the addition of 239 mM (20 mg/ml) methoxyamine hydrochloride in pyridine and incubation for 1.5 h at 30°C. Sequential addition of N-methyl-N-trimethylsilyl trifluoroacetamide followed with an incubation time of 30 min at 37 °C with thorough mixing between addition of solvents. The mixture was then transferred to a V-shaped amber glass chromatography vial and analyzed by GC-MS.

### GC-MS quantitation

An Agilent 7800B GC coupled to a 5977B MS detector was used for this study. GC-MS protocols were similar to those described previously [119], except a modified temperature gradient was used for GC with initial temperature at 60°C, held for 1 min, rising at 10°C/min to 325°C and held for 10 min. The electron ionization energy was set to 70 eV. Full scan mode (m/z, 50–800) was used for metabolomics analysis. The identification of metabolites was performed using the Automated Mass Spectral Deconvolution and Identification System (AMDIS) software matched to the FiehnLib metabolomics library (available through Agilent, Santa Clara, CA, USA) with retention time and fragmentation pattern matching [120–123]. Relative quantitation was performed using the software package DExSI (Data Extraction for Stable Isotope-labelled metabolites) (https://github.com/DExSI/). Relative abundance was corrected for recovery using the L-norvaline standard and adjusted to protein input.

### Amino acid uptake assay

Uptake assays were based on published procedures [124]. Cells grown for 24 h at 30°C in SDC or SC media, were diluted into 25 mls of fresh culture medium as described for lifespan assays. After incubation for the times indicated in Figure legends, 3 A600nm units of cells were collected on a membrane filter, 0.45 μm dia. (Millipore-Sigma cat. #HAW02500) and washed twice with 5 ml of assay solution (SDC medium lacking amino acids and NH4SO4, no pH adjustment) warmed to 30°C. All remaining steps were done at 30°C. Cells were removed by placing the filter into a 50 ml conical, plastic tube containing 1.235 ml of assay solution and vortexed 15 sec before removing the filter. Tubes were shaken at 15 second intervals for 2 min to acidify the medium and then 12.5 ul (1-1.5 x 10^6^ dpm) of a mix of radioisotope and non-radioactive substrate were diluted while vortexing into the assay solution. Three min later, 0.5 ml of this solution was filtered and washed twice with 5 ml cold water. Filters were transferred to 4 ml of Liquid Gold scintillation fluid and radioactivity was quantified in a liquid scintillation counter. Specific activities were calculated as described [124]. Specifically, a standard curve of mg protein/A600nm units verses time of cell growth (0, 2, 4 and 6 h) was made for myriocin-treated and untreated and used to adjust small differences in protein concentration/A600nm unit over time and the two treatments.

### Microscopy and protein trafficking assays

Mup1 localization and trafficking were analyzed by fluorescence microscopy and flow cytometry as previously described [69, 70] with minor modifications. Briefly, JTY240 (endogenously expressing Mup1-GFP and Vph1-MARS) and JMY1811 (endogenously expressing Mup1-pHluorin) cells were grown in Met-deficient SC medium and treated with 20 μg/ml methionine to stimulate Mup1 trafficking from the plasma membrane into the vacuole. Mup1-GFP localization in JTY240 cells was visualized over time using a DeltaVision Elite Imaging system (Olympus IX-71 inverted microscope; Olympus 100× oil objective (1.4 NA); DV Elite sCMOS camera, GE Healthcare). Mup1-pHluorin trafficking in JMY1811 cells was analyzed over time using the BD Accuri^TM^ C6 Plus Flow Cytometer by measuring the relative intensity (FITC channel) of pHluorin signal of 10,000 cells every sampling time point.

### Immunoblotting

JMY78 cells (with chromosomal Mup1-FLAG) were grown in SDC medium with or without Myr treatment. Total protein, from 5 A600nm equivalents of cells harvested from each time point, were precipitated by addition of 1 ml of ice-cold 10% trichloroacetic acid in TE buffer (50 mM Tris pH 8.0, 1 mM EDTA) and left on ice for 1 h. Precipitates were washed twice with ice-cold acetone, disrupted by sonication, dried by aspiration and vacuum centrifugation, resuspended in lysis buffer (50 mM Tris pH 7.5, 150 mM NaCl, 1 mM EDTA, 1% SDS), and vortexed vigorously with acid-washed glass beads (0.5 mm dia.). Proteins were diluted with equal volume of 2X sample buffer (150 mM Tris pH 6.8, 6 M urea, 6% SDS, 10% β-mercaptoethanol, 20% glycerol), separated by electrophoresis in 12% Bis-Tris polyacrylamide gels, and transferred onto a polyvinylidene difluoride membrane. The membrane was blocked with 3% bovine serum albumin in TBS-T buffer (50 mM Tris pH 7.5, 150 mM NaCl, 0.1% Tween-20) and incubated with anti-FLAG (Sigma, St. Louis, MO, cat# AB_262044, 1:1000 dilution) and anti-G6PDH (Sigma, St. Louis, MO, cat# AB_258454, 1:10000 dilution) primary antibodies followed with IRDye 680LT goat anti-mouse (LI-COR Biosciences, cat# 926-68020, 1:10000 dilution) and 800CW goat anti-rabbit (LI-COR Biosciences, cat# 926-32211, 1:10000 dilution) secondary antibodies. Fluorescence was visualized using the Odyssey CLx imaging system and quantified using Image Studio Lite (LI-COR Biosciences).

## Author contributions

Project Conception: RCD, NLH and JAM; Execution: NLH, JKAM, LEAY, KL, RCS and RCD. Data analysis: RCS, JAM, NLH and RCD. Manuscript: Writing: RCD and JAM; Review and Critique: NLH, JKAM, LEAY, KL, RCS.

## Acknowledgements

We thank Dr. Mathew Gentry for access to instrumentation and disposable supplies.

## Supplemental Data

**S1 Figure. Myr treatment increases longevity in buffered SDC medium**.

The lifespan of auxotrophic or prototrophic BY4741 cells was determined in SDC medium buffered with citrate-phosphate buffer and 1.50 μmol/L (600 ng/ml) of myriocin as previously described (Liu J, Huang X, Withers BR, Blalock E, Liu K, Dickson RC. Reducing Sphingolipid Synthesis Orchestrates Global Changes to Extend Yeast Lifespan. Aging Cell. 2013;12:833-41).

**S2 Figure. Initial cell cycle progression.**

To examine how synchronized stationary phase cells are when inoculated into fresh SDC culture medium and to determine when they complete their first cell division cycle, we measured their budding behavior by using the Budding Index assay [125].

**S3 Figure. Myr treatment reduces Mup1 activity in the PM.**

Flow cytometry analysis like that shown in Figure 5C, except cells were cultured in SDC medium, which contains 537 μmol/L methionine. As shown, flow cytometry results of cells cultured in this manner results in low surface expression of Mup1-pHluorin that is below the limit of useful detection for this assay. Thus, for culturing cells in SDC media, Mup1-FLAG was used since immunodetection of FLAG epitope in yeast lysates is more sensitive for measuring Mup1 expression (Figure 5E-F).

**S1 Table. Free amino acids pools.**

Free amino acid pools were measured in auxotrophic DBY747 cells (require Ura, Leu, Trp and His) grown in SDC medium buffered with succinate (Liu J, Huang X, Withers BR, Blalock E, Liu K, Dickson RC. Reducing Sphingolipid Synthesis Orchestrates Global Changes to Extend Yeast Lifespan. Aging Cell. 2013;12:833-41). Cells were grown from 0.005 to 2 A600nm units/ml at 30°C without and with 0.75 μmol/L (300 ng/ml) of myriocin. Amino acids were extracted by using the heat extraction procedure described in Materials and Methods and quantified by using a Hitachi L-8800A amino acid analyzer. The values for Asp are a combination of Asp and Asn, and for Glu they are combination of Glu and Gln. This occurs because the procedure for preparing samples for injection into the amino analyzer cause deamination of Asn and Gln. Both of these amino acids have small intracellular pools compared to Asp and Glu (see Figure 2). AUC Auxo-No Myr vs Auxo-Plus Myr (95% CI 14.89 to 15.97 vs 20.46 to 21.90). AUC Proto-No Myr vs Proto-Plus Myr (95% CI 25.54 – 26.66 vs 27.82 – 33.65).

