## Supplementary figures and images for "Enhancing lifespan of budding yeast by pharmacological lowering of amino acid pools"

### Supplemental Figure S1

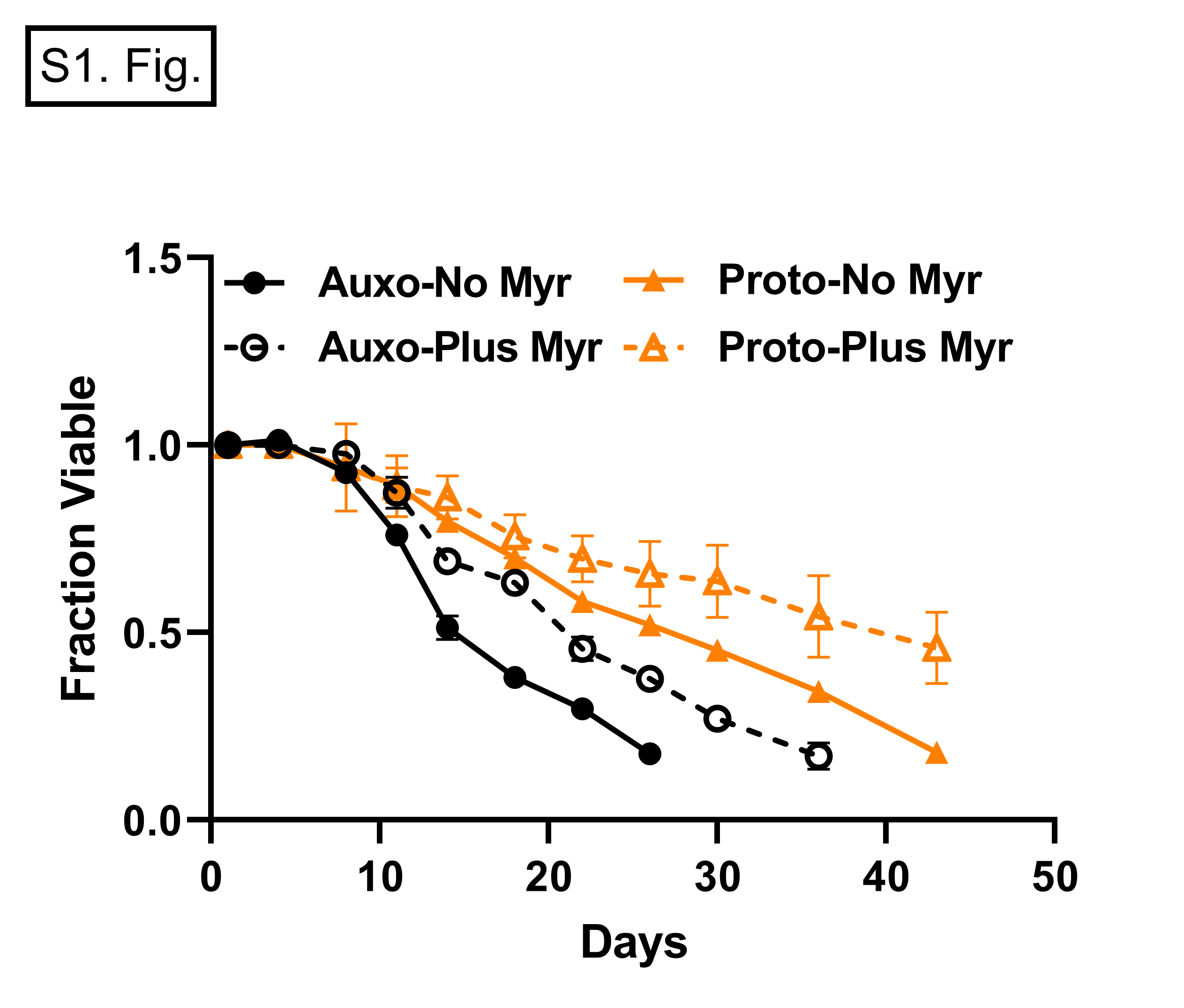

### Supplemental Figure S2

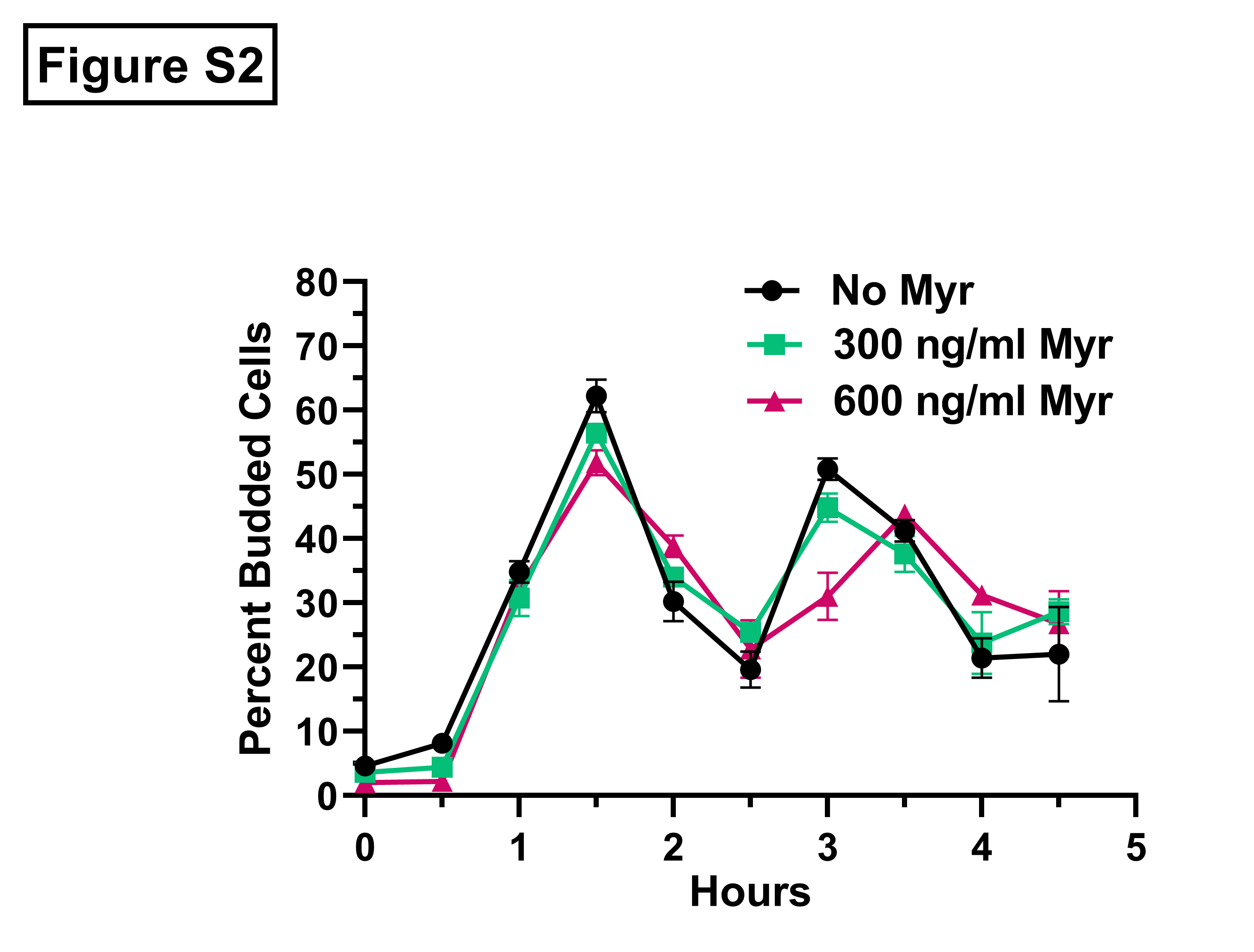

### Supplemental Figure S3

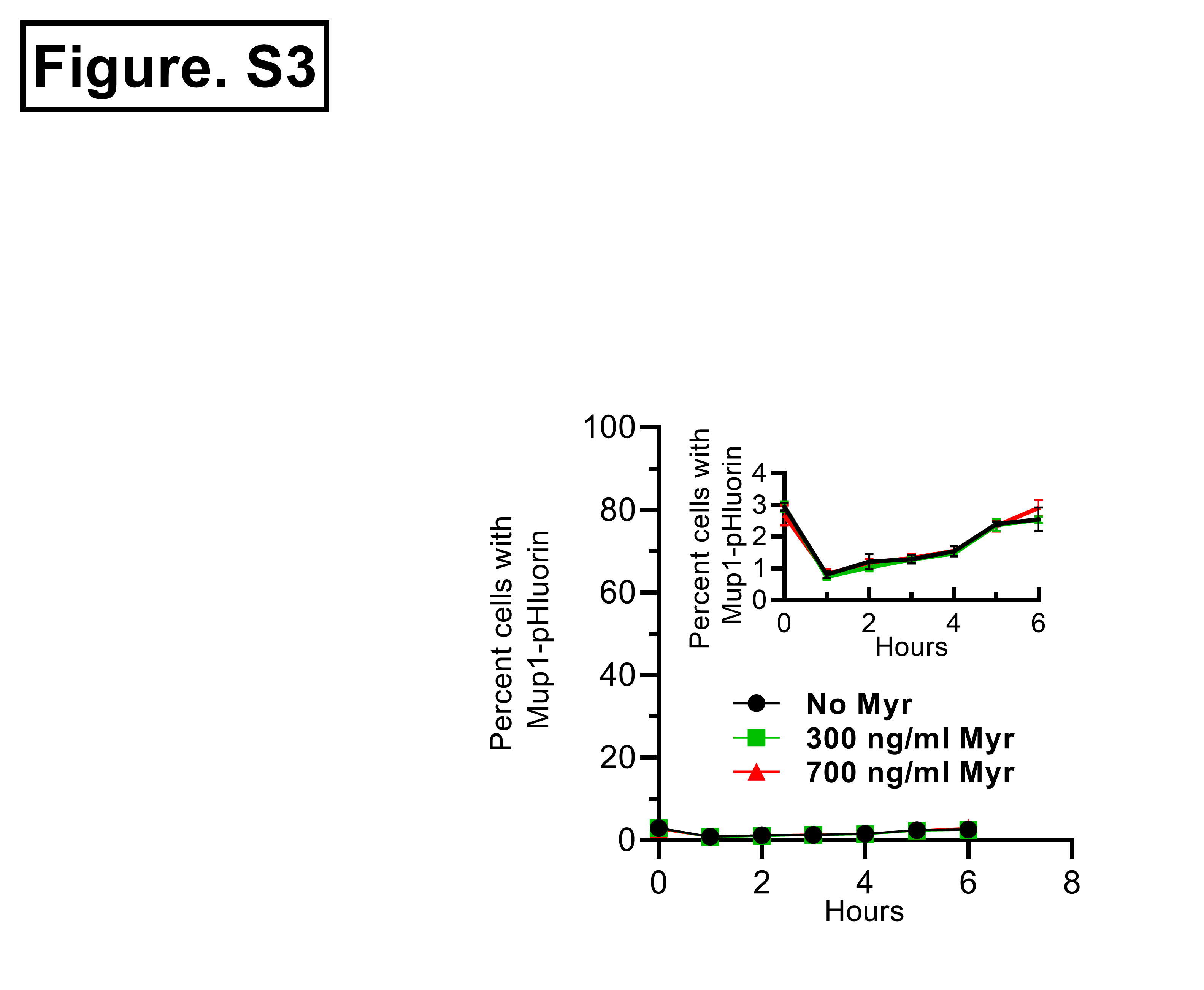
